# A *Varroa destructor* protein atlas reveals molecular underpinnings of developmental transitions and sexual differentiation

**DOI:** 10.1101/144808

**Authors:** Alison McAfee, Queenie WT Chan, Jay Evans, Leonard J Foster

## Abstract

*Varroa destructor* is the most economically damaging honey bee pest, weakening colonies by simultaneously parasitizing bees and transmitting harmful viruses. Despite these impacts on honey bee health, surprisingly little is known about its fundamental molecular biology. Here we present a *Varroa* protein atlas crossing all major developmental stages (egg, protonymph, deutonymph and adult) for both male and female mites as a web-based interactive tool (http://foster.nce.ubc.ca/varroa/index.html). By intensity-based label-free quantitation, 1,433 proteins were differentially expressed across developmental stages, including two distinct viral polyproteins. Enzymes for processing carbohydrates and amino acids were among many of these differences as well as proteins involved in cuticle formation. Lipid transport involving vitellogenin was the most significantly enriched biological process in the foundress (reproductive female) and young mites. In addition, we found that 101 proteins were sexually regulated and functional enrichment analysis suggests that chromatin remodeling may be a key feature of sex determination. In a proteogenomic effort, we identified 519 protein-coding regions (169 of which were differentially expressed) supported by 1,464 peptides which were previously unannotated. Since this is a recurring trend with annotating genomes of non-model species, we analyzed their amino acid and nucleotide composition as well as their orthology to other species to suggest reasons why they may have been missed initially. Overall, this work provides a first-of-its-kind interrogation of the patterns of gene expression that govern the *Varroa* life cycle and the tools we have developed will support further research on this threatening honey bee pest.

## Introduction

The *Varroa destructor* mite is the most devastating pest for Western honey bees *(Apis mellifera)* (1–3). This obligate parasite feeds on honey bee hemolymph (blood), simultaneously weakening its host, suppressing the innate immune system and transmitting debilitating viruses (see Rosenkranz *et al.* (4) for a comprehensive review on *Varroa* biology). *Varroa’*s natural host is the Eastern honey bee *(A. cerana),* and millions of years of co-evolution has led *A. cerana* to develop various tolerance mechanisms, thereby minimizing the mite’s negative impact on these colonies (5–7). However, in the mid-1900s the mite jumped hosts to *A. mellifera* – the bee species that is most commonly used for active crop pollination today – which is less effective at defending itself (4, 6). Managed *A. mellifera* colonies infested with *Varroa* have shorter lifespans than uninfested colonies unless they are actively treated with miticides (8, 9), causing serious negative economic impacts (10–12).

Despite being responsible for significant colony losses, very little is known about the molecular biology of the *Varroa* mite. Since the egg, protonymph and deutonymph life stages (**Figure 1**) only exist when the foundress mite (reproductive female) is actively reproducing within capped honey bee brood comb (4), they are seldom observed and are tedious to sample. Furthermore, male mites (even as adults) die soon after the adult honey bee emerges so even though they are obviously important factors in mite reproduction, our knowledge of their basic molecular biology is extremely limited. Research on *Varroa* has focused on its role as a vector for viruses (13–18), their response to pheromone cues (19–21), attempts to control it via RNAi (22–24) and host shifts (25). At the time of writing, there have only been two previous *Varroa* proteomic investigations, one of which focused on viral proteins (15) and the other identifying fewer than 700 proteins within one developmental stage (26). Global gene expression changes associated with developmental transitions and sexual differentiation are yet unknown.

**Figure 1.**
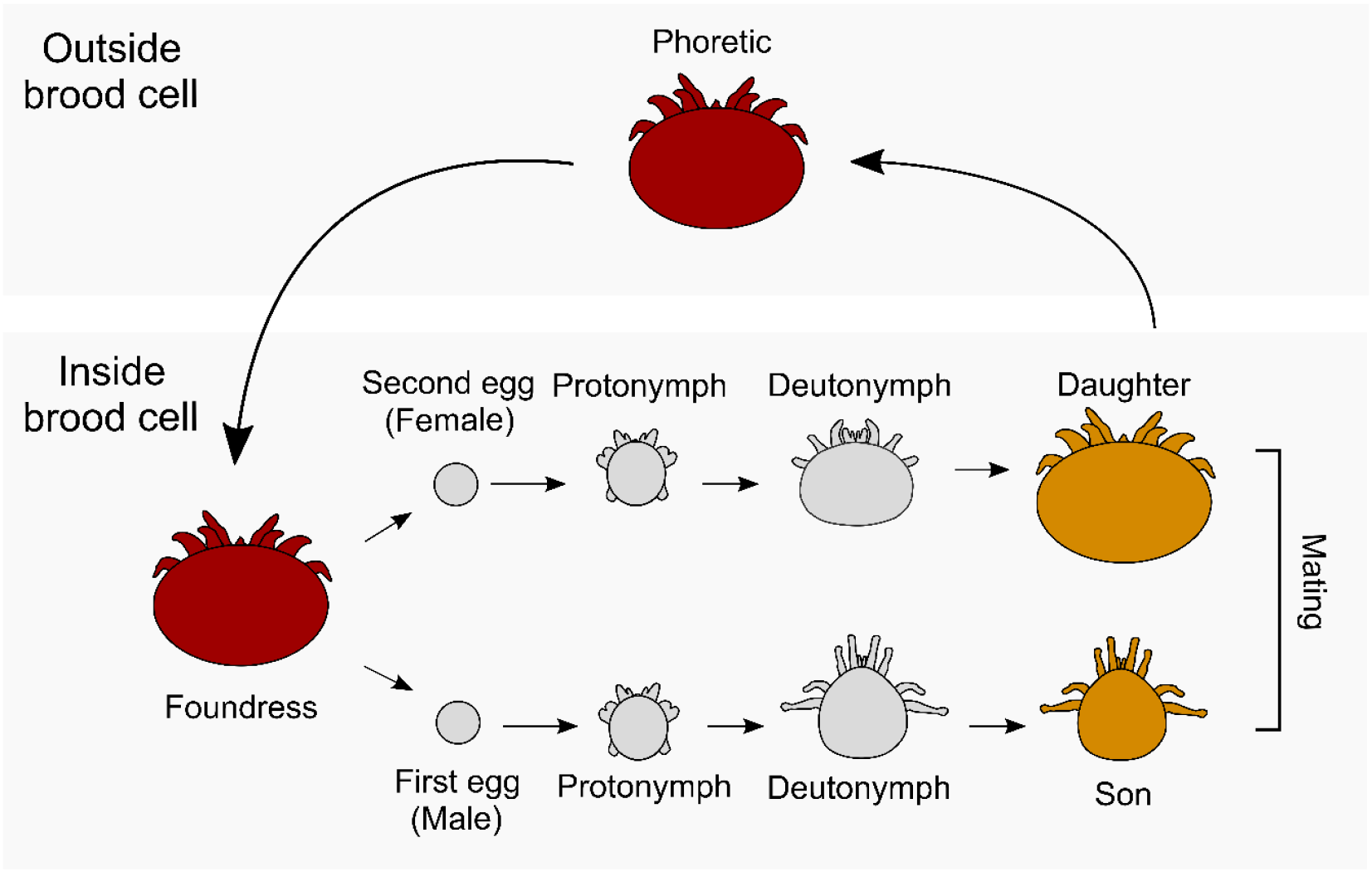
Schematic representation of the mite life cycle. All stages were included in this study (n = 3 for all) except the phoretic stage. For egg and protonymph stages, males and females are visually indistinguishable so for these stages, sexes were pooled. Colours indicate melanisation of the cuticle and sizes are proportional.

The *Varroa* genome was first sequenced in 2010 (1) and was accompanied by a provisional gene annotation that will be updated shortly^1^. Gene annotations are living databases and, particularly with newly sequenced species, they undergo continuous refinement as more ‘omic data becomes available. Unfortunately, the more evolutionarily distant a species is from well-annotated species typically used for orthology delineation and gene prediction training sets, the less accurate the predictions become. Such is the case for *Varroa.* Proteogenomics (27, 28) can help overcome this problem by sequencing the expressed protein regions in a relatively unbiased survey of the genomic landscape. Since gene expression is dynamic throughout an organism’s life cycle, high resolution ‘omics data that crosses developmental stages and sexes is very well-suited for this purpose.

Investigating global gene expression profiles throughout development of both sexes simultaneously provides a foundational understanding of *Varroa* biology and creates an opportunity to improve upon existing gene annotations. We present here the first *Varroa* proteome crossing all major developmental stages (egg, protonymph, deutonymph, adult) of both males and females, where distinguishable (**Figure 1**). Through a proteogenomics effort, we identified 519 new protein-coding regions. We also analyze the chemical properties of these sequences and their sequence similarity to other organisms to suggest reasons why under-annotation continues to be a problem. We identified 3,102 proteins overall, nearly half (1,433) of which were significantly differentially expressed through development, and 101 of which were differentially expressed between sexes. Functional enrichment suggested that carbohydrate and amino acid metabolism underpin developmental transitions, so we investigated proteins involved in glycolysis and the Krebs cycle in detail. Cuticle formation is clearly a process associated with mite aging, and closer analysis suggests the mites utilize different chitin structural proteins as they mature. In addition, chromatin remodelling and positive regulation of transcription may be key factors in sexual differentiation. Building on our previous honey bee protein atlas (29), we provide a web-based interactive platform [http://foster.nce.ubc.ca/varroa/index.html] where researchers can query proteins for visual displays of expression patterns, enabling further hypothesis generation and maximizing the utility of this information for the scientific community.

## Experimental Procedures

### Sample collection

*Varroa* mite families were collected from a single *A. mellifera* colony in the fall of 2016 in Vancouver, Canada. In a large-scale population genomics study, the authors found that the genetic variation of *Varroa* within colonies accounted for by far the largest fraction of genetic variation compared to between colonies and between apiaries (30); therefore, sampling mites from a single colony was sufficient. Eggs, foundresses, adult daughters and adult sons were transferred directly to microfuge tubes using a soft paintbrush, whereas protonymphs and deutonymphs were transferred to a petri dish and sorted under a dissecting microscope according to the identification guides available at http://idtools.org/id/mites/beemites and http://extension.msstate.edu/publications (publication number: P2826) *via* the University of Michigan and the Mississippi State University, respectively. Approximately 50 individuals were pooled for each replicate (7 developmental stages, n = 3 for each stage). All samples were immediately frozen at -72 °C until protein extraction.

### Protein preparation

Protein was extracted by homogenizing each mite stage with ceramic beads as previously described (31). Clarified lysate was precipitated overnight with 4 volumes of 100% ice cold acetone and the pellet was washed twice with ice cold 80% acetone. After allowing residual acetone to evaporate (~15 min) the protein pellet was solubilized in urea buffer (6M urea, 2M thiourea in 10 mM HEPES, pH 8) and ~30 μg (determined via the Bradford Assay) was reduced, alkylated and digested as previously described (32). Peptides were acidified (1 volume 1% TFA), desalted on a high capacity C18 STAGE tip (33), solubilized in Buffer A (0.1% formic acid) and quantified in technical triplicate using a peptide fluorimetric assay (Pierce).

### Data acquisition

Two μg of peptides per sample were analyzed on an EasynLC-1000 chromatography system (Thermo) coupled to a Bruker Impact II Q-TOF mass spectrometer. The LC C18 columns included a fritted trap column and pulled-tip, 50 cm analytical column produced and packed inhouse (34, 35). Peptides were separated using a 165 min linear gradient of increasing Buffer B as specified in the LCParms.txt file embedded within the Bruker data folders (available at www.proteomexchange.org, accession: PXD006072). Buffers A and B were 0.1% formic acid and 0.1% formic acid, 80% acetonitrile, respectively. The instrument was set to the same parameters as described in our previous publication under “Analysis of PTMs” (34), except the scanned mass range was 200-2,000 m/z, the top 20 precursors were fragmented at a 5 Hz spectral rate and the lower precursor intensity threshold was 300 counts.

### Mass spectrometry data analysis

#### Proteogenomics

For the proteogenomics analysis, the *Varroa* fragment spectra were searched against a six-frame translation of the publicly available *Varroa* genome sequence (PRJNA33465) using MaxQuant (v.1.5.3.30) to identify new protein-coding regions (minimum ORF length was set to 100 amino acids). All viruses known to infect *A. mellifera* and *Varroa* were also included in the database. Honey bee proteins were not included after a follow-up sequence similarity analysis indicated that only 5 of the proteins identified in this search matched to bees. MaxQuant search settings were the same as previously described (31). The peptide (scores, modifications, precursor mass and m/z) and protein (protein groups, accessions, number of assigned peptides, unique peptides and *%* coverage) identification information contained within the main MaxQuant output files (summary.txt, peptides.txt, proteinGroups.txt, parameters.txt) and the protein database (165,951 entries) are available at PXD006072.

Peptides identified in the six-frame translation search but which were not present in the canonical protein database were used as anchors to retrieve the corresponding open reading frames from the genome using a simple Perl script. This yielded 524 new protein-coding sequences. Of these, 301 were flanked by two or more peptides spanning at least 50 amino acid residues. We used a two-way ANOVA (factors: amino acid and new/known sequence origin) to compare amino acid composition between this set of 301 new protein-coding sequences and 902 sequences bounded by known peptides that were identified in the same six-frame translation search. Next, we used the same approach to compare nucleotide positions within codons (factors: nucleotide position and sequence origin). We also compared the AT frequency of the new coding regions, known coding regions and the *in silico* 1 kb fragmented genome (n = 384,129) using a 1-way ANOVA (3 levels) with a Tukey HSD post-hoc test. We have included the Perl script modules used in these analyses as **Supplemental File 1**.

To survey these proteins for orthology with other species and to retrieve GO terms, we performed Blast2GO (v4.0) using default parameters. We reasoned that these sequences might have been missed in the *Varroa* annotation effort if they only share sequence similarity to evolutionarily distant species; therefore, we queried them against the non-redundant protein collection with no taxonomic restrictions. Five sequences showed significant homology to honey bee sequences and were removed from the list of new protein coding fragments, leaving 519 in total.

#### Protein quantitation

We searched the mass spectrometry data using the same parameters as above, except label-free quantitation (LFQ) was enabled and a composite protein database was used which included all proteins in the most recent *Varroa* gene annotation^1^ (the final protein database is included at the ProteomeXchange accession below), the 519 protein coding fragments identified above, all virus sequences known to infect *A. mellifera* or *Varroa* and all proteins contained within the *A. mellifera* OGSv3.2 annotation. Since *A. mellifera* biological material is *Varroa’s* sole food source, we expected to find a substantial number of honey bee proteins within our samples. The final database totaled 32,110 entries and is available at PXD006072, along with the MaxQuant peptide and protein identification information as described under “*Proteogenomics*.” Honey bee proteins include an “Amel” tag in the accession, new protein-coding regions from the six-frame translation include a “True” or “False” tag in the accession (indicating the DNA template strand relative to the indicated contig), virus sequences are represented by a single gi number or Uniprot identifier and all other sequences (excluding contaminants and reverse hits) belong to *Varroa.* LFQ parameters were the same as previously described (31).

#### Experimental design and statistical rationale

All seven developmental stages were collected in biological triplicate with approximately 50 individuals pooled to create each replicate. Since each stage is a pool of many individuals, even a relatively low replication of n = 3 represents a large sample of the population. Only proteins with 6 or more observations (out of 21) were included in differential expression analysis across developmental stages. For the differential expression across sexes, this was relaxed to 3 or more observations to avoid excluding proteins that are stage- and sex-specific. Visual inspection of log-transformed LFQ intensity histograms confirmed the data for each replicate was distributed normally prior to analyzing with an ANOVA. Differential expression analysis across developmental stages was performed as previously described (31) except Perseus v1.5.6.0 was used, missing values were not imputed and the ANOVA (one factor, 7 levels) p-values were Benjamini Hochberg-corrected at 5% FDR. For the analysis of sexually regulated genes, the female (deutonymph, adult daughter and foundress) and male (deutonymph, adult son) samples were pooled as n = 9 and n = 6, respectively. Since there was a large amount of presence/absence data between males and females (potentially the most interesting differences), missing values were imputed from a normal distribution (Perseus parameters: width = 0.3; downshift = 1.5) as previously described (36). A t-test was then performed and subjected to the Benjamini Hochberg correction at 5% FDR. All hierarchical clustering analyses were performed in Perseus using average Euclidian distance (300 clusters, maximum 10 clusters).

### Functional enrichment analysis

We performed functional enrichment analysis on two sets of proteins: 1) *Varroa* proteins that were differentially expressed through development and 2) *Varroa* proteins that were differentially expressed between sexes. For all protein sets, we retrieved GO terms using Blast2GO (v4.0) with default parameters, first searching against all arthropods, then sequences with missing GO terms were searched again against the entire non-redundant protein collection. GO terms were exported after running the GO-Slim function. We then performed a gene score resampling (GSR) analysis with ErmineJ v3.0.2 (37), using log-transformed q values (from the previous differential expression analysis) for “protein score.” We considered a GO term significantly enriched if the Benjamini Hochberg-corrected GSR p value was less than 0.10.

### *Building the* Varroa *protein atlas*

The web-based interactive *Varroa* protein atlas was built using the framework previously described for the honey bee protein atlas (29).

## Results

### *The new* Varroa *gene set has dramatically improved accuracy over the first draft*

Procuring an accurate protein database is critically important for proteomics applications. The first *Varroa* draft gene set was published in 2010 (1) along with the initial genome sequence (ADDG00000000.1); however, a new genome build was just released (ADDG00000000.2) with annotation refinement efforts underway.^1^ A new gene set will soon to be released, and we have made the new protein database provisionally available through ProteomeXchange (PXD006072). To test the accuracy of the new gene set compared to the first draft, we searched our complete *Varroa* proteomics data against both versions and found that greater than 2-fold more unique peptides were identified using the refined annotation (**Figure 2A**). Overall, we identified nearly 20,000 unique peptides corresponding to 3,102 protein groups at 1% peptide and protein FDR (**Figure 2B**) representing the first global survey of *Varroa* protein expression. To maximize the utility of this information for researchers, we incorporated the quantified proteins into an interactive *Varroa* protein atlas (http://foster.nce.ubc.ca/varroa/index.html). The atlas features a searchable database of the quantified proteins as well as a visual and numerical display of their relative expression in different developmental stages (**Figure 3**).

**Figure 2.**
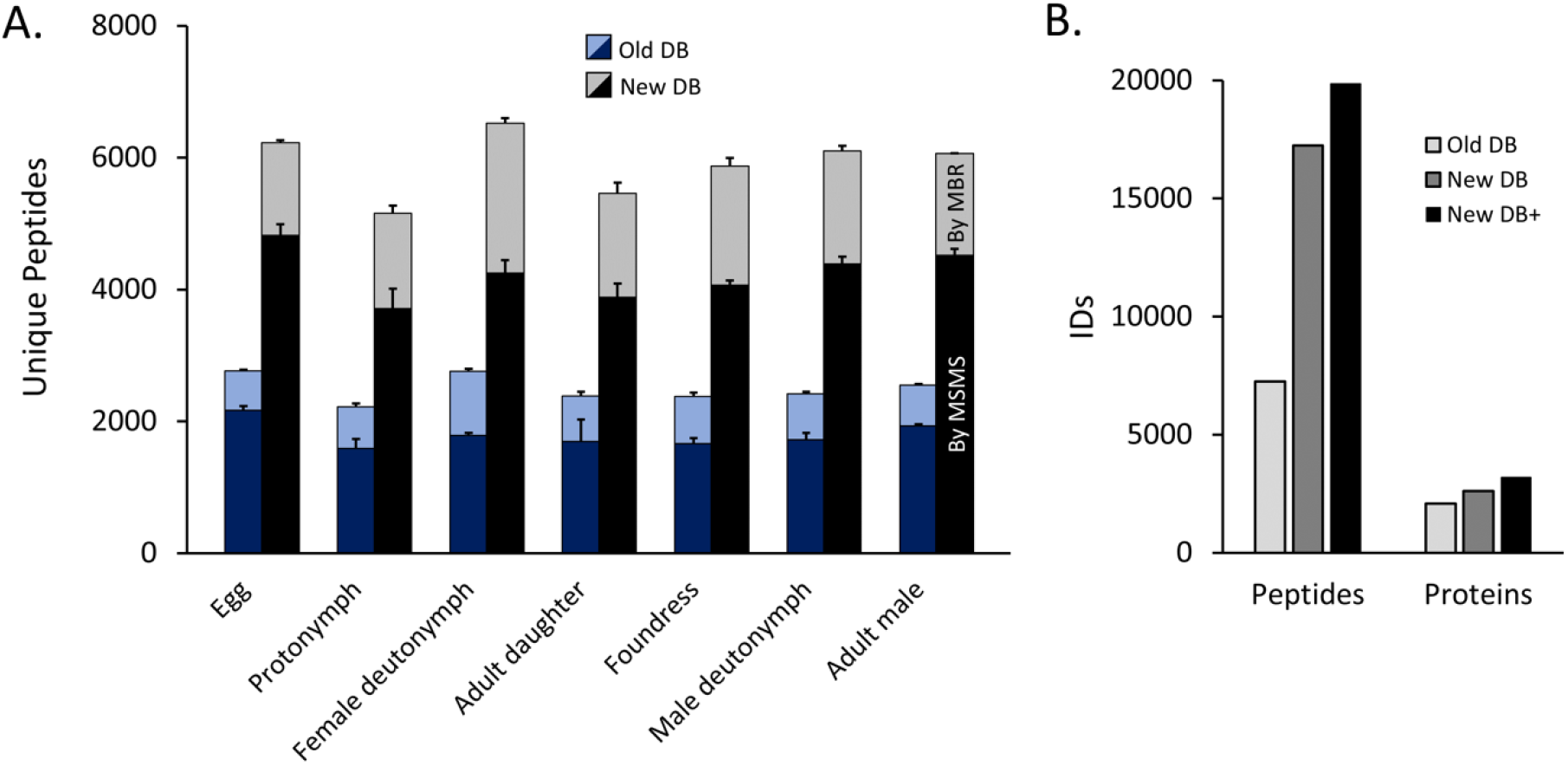
Overall peptide and protein identifications. MSMS data was searched against the initial draft *Varroa* gene annotation (Old DB) and the most recent updated annotation (New DB). The data included biological triplicates of each developmental stage and all protein databases also included NCBI *Varroa* sequences and all viruses known to infect honey bees and *Varroa.* A) light stacks represent peptide identifications via match between runs (MBR) and dark stacks represent identifications via MSMS matching. Error bars are standard deviation. B) Cumulative identifications. New DB+ refers to the newest annotation plus all honey bee proteins and new fragments identified by proteogenomics.

**Figure 3.**
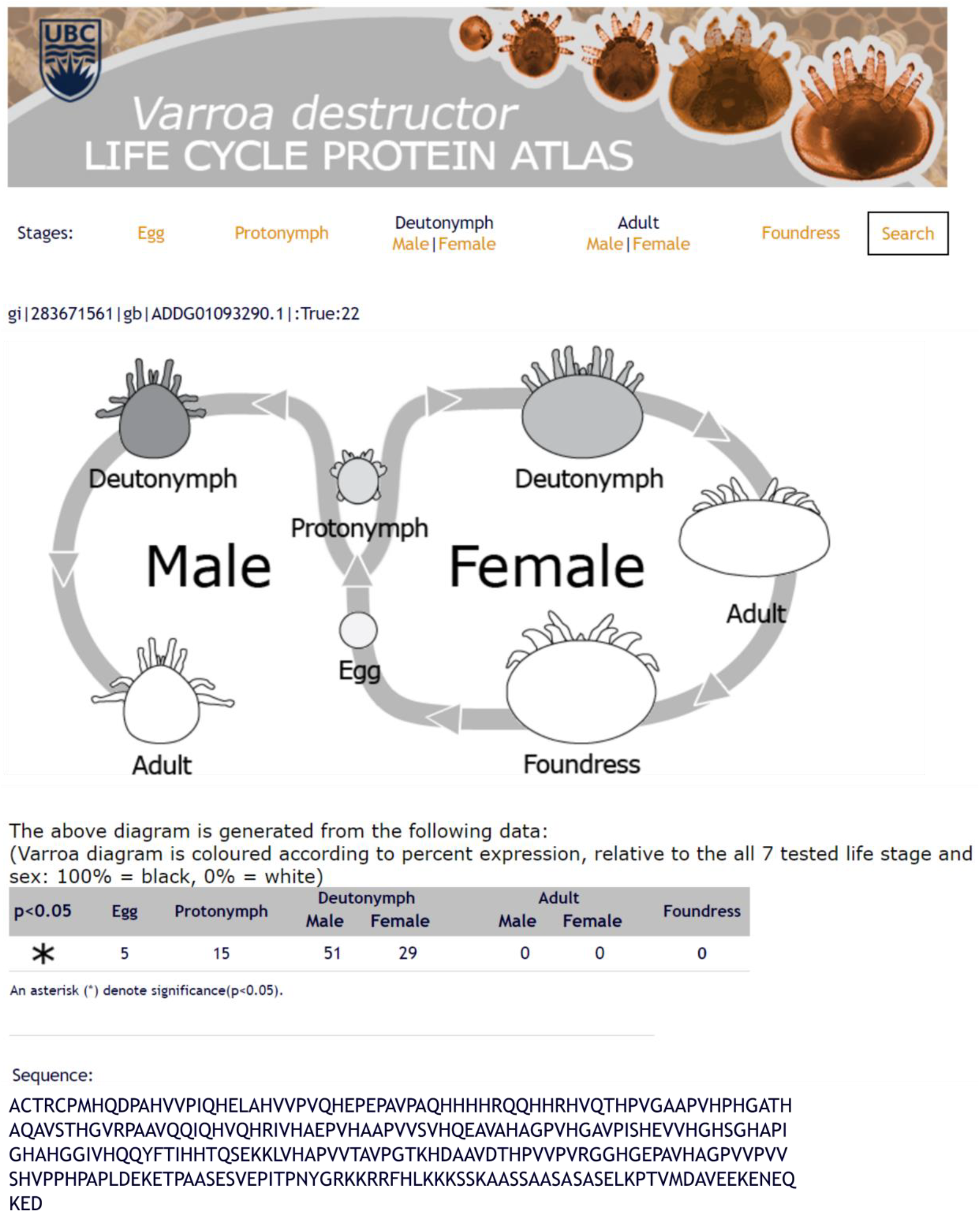
Example page of the web-based. Varroa destructor *protein atlas.* The atlas was constructed using the framework described for the honey bee protein atlas (29). Shading of the cartoon mites indicates relative expression and an asterisk indicates that this protein was significantly differentially expressed according to developmental stage. Website: http://foster.nce.ubc.ca/varroa/index.html.

### Proteogenomics identifies unannotated regions

Despite a dramatic improvement in accuracy over the initial draft annotation, the current annotation could likely be further improved through proteogenomics. We searched the MSMS data against a six-frame genome translation database and identified 524 protein-coding regions at 1% FDR (see **Supplementary Table 1** for protein and peptide sequences) which were absent from the current annotation. Furthermore, 169 of these protein groups were differentially expressed through development (**Figure 4A**). This is in line with improvements we have discovered previously in *A. mellifera,* another non-model organism (34); however, since missed genes appear to be a common problem in genome annotation, we sought to investigate the root cause of failing to locate these sequences in the first place.

Gene prediction algorithms often use training gene sets from well-annotated species with similar genomic properties to help define genes in the newly sequenced target species (27). We hypothesized that one reason why an algorithm might fail to identify expressed sequences is if they occur in regions with significantly different AT content or codon bias (indeed, this is precisely what happened during the *A. mellifera* annotation (38)), so we compared these properties between the newly identified protein coding regions and the previously known coding regions identified in the same six-frame translation search. We found that the newly identified regions had the same AT content as the previously known regions, which were both significantly different from the genomic average (**Figure 4B**). While this lends additional confidence that the new regions are expressed, it does not explain why they were missed. Furthermore, the amino acid composition (**Figure 4C**) and nucleotide positional codon bias (**Figure 4D**) was the same between the new and known coding regions.

Since some algorithms rely on homology evidence to support annotations, one reason sequences may not be annotated is if they do not have known orthologs. We used Blast2GO to identify potential orthologs and found that nearly 72% (377) of the sequences had significant similarity (e-value cut-off: 1E-5) to at least one sequence in the non-redundant NCBI protein database (**Figure 4E**). Of those, the majority (85%) matched to sequences from other members of phylum Arthropoda but Chordata, Nematoda, Mollusca and Annelida were also present. Importantly, only nine sequences significantly matched well-annotated species *(Homo sapiens, Mus musculus, Drosophila melanogaster* and *Caenorhabditis elegans)* and 148 (28%) had no significant sequence similarity to any species. In addition, 5 sequences were highly similar to known honey bee sequences, suggesting these are likely the result of DNA contamination within the *Varroa* sample used for DNA sequencing. This is not surprising since honey bee tissue is the mite’s sole food source, so some contamination of this nature is expected. We removed these sequences since we include all honey bee proteins in our search database regardless in order to account for abundant honey bee proteins consumed by *Varroa.* However, we cannot discount the possibility that these 5 protein fragments are truly components of *Varroa* genes that just happen to be highly similar to honey bees or which are the result of horizontal gene transfer. All other fragments identified through proteogenomics were added to the protein database and utilized in subsequent analyses.

**Figure 4.**
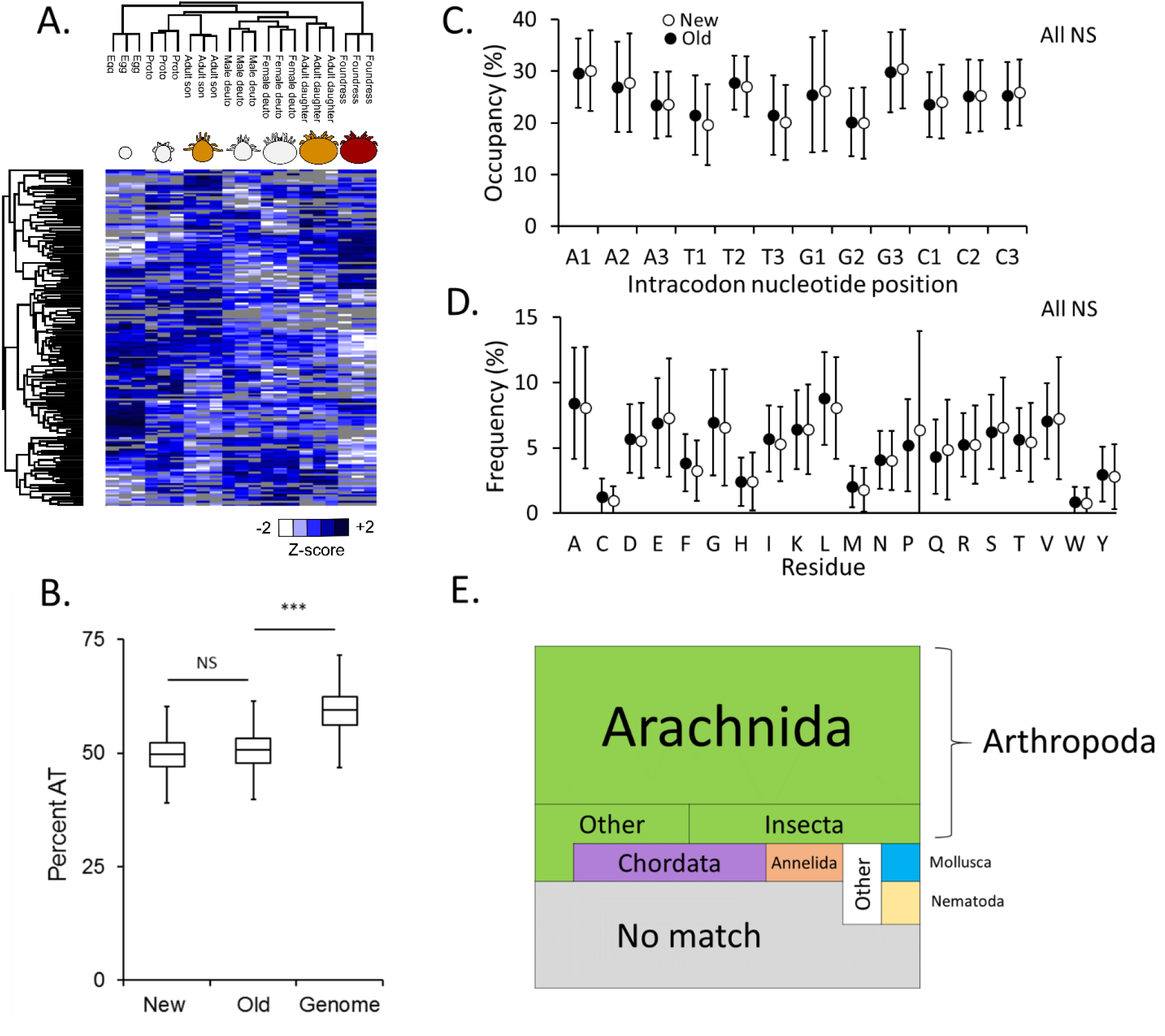
A search against a six-frame genome translation database identifies new protein-coding fragments. A) New protein-coding fragments which are differentially regulated across development. Grey tiles represent missing data. Hierarchical clustering was performed in Perseus using average Euclidian distance (300 clusters, maximum 10 clusters). Statistics were performed using an ANOVA (Benjamini Hochberg-corrected FDR = 5%). B) Comparison of the AT composition between newly identified sequences, previously known sequences (old) and the *in silico* fragmented genome. Statistics were performed using a one-way ANOVA (3 levels) and a Tukey HSD post-hoc test. NS: not significant. ***: p < 0.0001. Boxes depict the interquartile range (IQR) and whiskers span 1.5*IQR. There are no significant differences in mean amino acid composition (C) or intracodon nucleotide position (D) of new protein coding regions compared to old (one way ANOVA). Error bars represent standard deviation. E) BLAST sequence alignment summary of the new protein coding regions for major (> 1% frequency) taxa. Area is proportional to frequency.

### Vitellogenin, carbohydrate metabolism and chitin expression underpin developmental transitions

Of the 3,102 proteins identified, 1,433 were significantly differentially expressed across developmental stages (**Figure 5A**; **Table 1**). As a quality control method, we specifically analyzed vitellogenin (an evolutionarily conserved yolk protein) expression since this is one of the only proteins where the developmental patterns of expression are known (39). We expected to see high levels of vitellogenin-1 and vitellogenin-2 in the foundress and egg, with quantities decreasing approaching adulthood (39) and indeed, this is what was observed (**Figure 5B**). Interestingly, some of the novel peptides identified in our proteogenomic effort mapped back to protein fragments with significant sequence similarity to vitellogenin, and upon closer inspection we found that some of these peptides are simply non-synonymous single nucleotide sequence variants of this well-known gene. However, we also identified novel protein fragments with significant similarity to vitellogenin that did not physically overlap with the known vitellogenin genes (**Figure 5C**). Like vitellogenin-1 and 2, the highest protein abundance for these novel sequences was in the egg. Furthermore, they group into two clusters of expressed fragments (one two-fragment cluster and one four-fragment cluster) closely linked on two different contigs, suggesting that the fragments form exons of two different genes (**Figure 5D**) and clearly illustrate how mass spectrometry data can aid in gene predictions.

**Figure 5.**
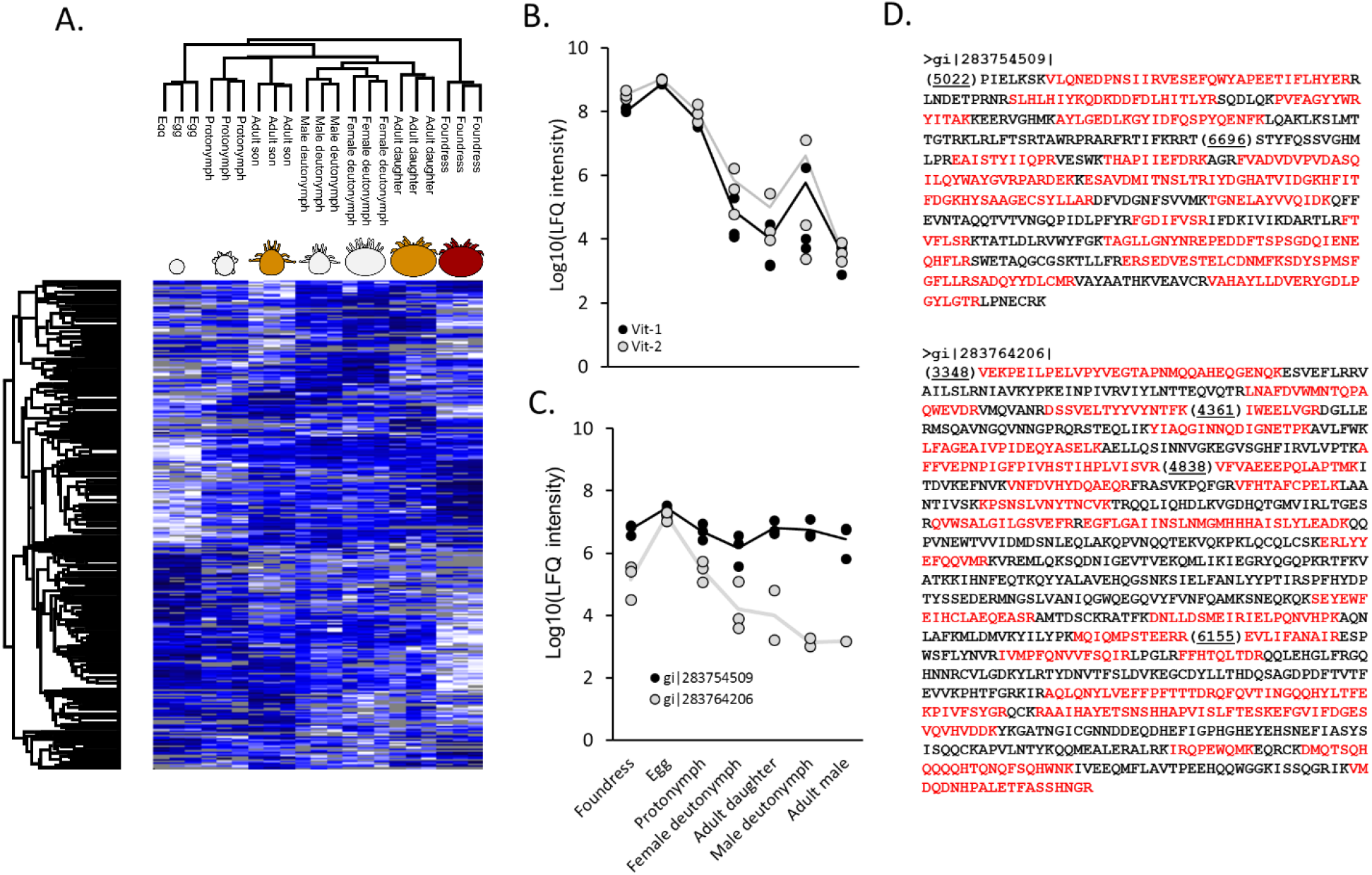
Analysis of vitellogenin expression. A) Heatmap showing significantly differentially expressed proteins across developmental stages (ANOVA; Benjamini Hochberg-corrected FDR = 5%).Grey tiles indicate missing data. Proto and deuto refer to protonymph and deutonymph, respectively. B) Vitellogenin (Vit)-1 and Vitellogenin-2 protein expression across developmental stages. C) Expression of new protein fragments showing significant vitellogenin homology. D) New vitellogenin protein fragment sequences. Observed peptides are red, the fasta header indicates the contig number and bracketed numbers indicate the nucleotide start position of each fragment within the contig. Both protein sequences were coded on the reverse contig strand.

**Table 1.**
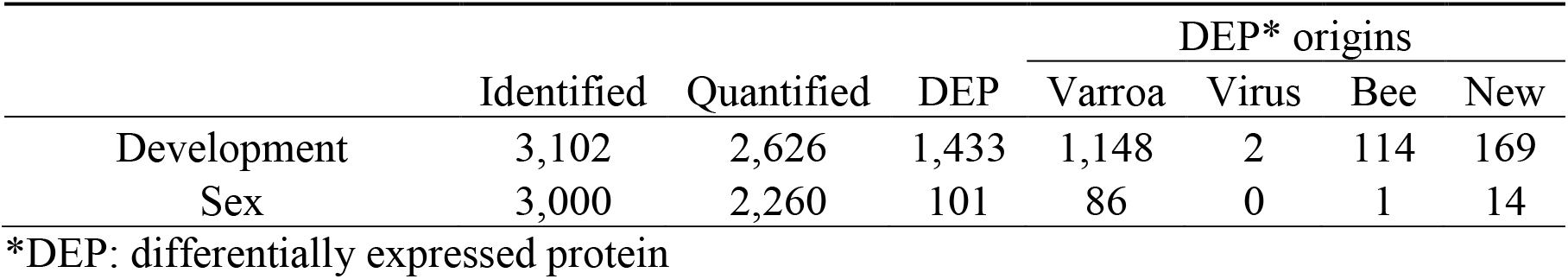
Summary of protein identifications

|  | Identified | Quantified | DEP | DEP* origins |  |  |  |
| --- | --- | --- | --- | --- | --- | --- | --- |
|  |  |  |  | Varroa | Virus | Bee | New |
| Development | 3,102 | 2,626 | 1,433 | 1,148 | 2 | 114 | 169 |
| Sex | 3,000 | 2,260 | 101 | 86 | 0 | 1 | 14 |

\*DEP: differentially expressed protein

To gain a better understanding of the cellular processes underlying proteins that were differentially expressed through development, we performed an enrichment analysis by gene score resampling (GSR) and found, not surprisingly, that lipid localization and lipid transport were among the most significantly enriched (**Table 2**, **Supplementary Table 2**), driven largely by vitellogenin expression. Many processes involved in aerobic respiration were also significantly enriched, including GO terms linked to glycolysis (GO:0006090, GO:0006096) and the citric acid cycle (GO:006099, GO:0072350). To investigate these metabolic processes further, we analyzed how the abundances of core glycolysis and citric acid cycle enzymes varied with development (**Figure 6A**). Most enzymes (16/20) were significantly differentially expressed and only two (phosphoglyceromutase and succinyl CoA synthetase) were not quantifiable. Several enzymes appear to have multiple isoforms, based on BLAST search results, some of which are not co-expressed (e.g. for hexokinase, α-ketoglutarate dehydrogenase, aconitase, isocitrate dehydrogenase and malate dehydrogenase). Overall, the foundress mite has the highest levels of most enzymes, and when this is not the case it is largely due to age-specific isoform expression. Relative expression levels for each protein can be found in **Supplementary Table 3**.

**Table 2.** GO terms significantly enriched in developmental stages

| Category | Name | ID | # Genes | Corrected p value* |
| --- | --- | --- | --- | --- |
| Glycolysis & TCA | aerobic respiration | GO:0009060 | 15 | 0.0777 |
|  | carbohydrate metabolic process | GO:0005975 | 62 | 0.0556 |
|  | dicarboxylic acid metabolic process | GO:0043648 | 10 | 0.0551 |
|  | tricarboxylic acid cycle | GO:0006099 | 13 | 0.0509 |
|  | glycolytic process | GO:0006096 | 10 | 0.0526 |
|  | cellular respiration | GO:0045333 | 22 | 0.0531 |
|  | monocarboxylic acid metabolic process | GO:0032787 | 36 | 0.0358 |
|  | energy derivation by oxidation of organic compounds | GO:0015980 | 24 | 0.0343 |
|  | tricarboxylic acid metabolic process | GO:0072350 | 14 | 0.0288 |
|  | pyruvate metabolic process | GO:0006090 | 16 | 0.0179 |
|  | generation of precursor metabolites & energy | GO:0006091 | 36 | 4.5E-10 |
|  | ADP metabolic process | GO:0046031 | 11 | 0.0544 |
|  | ATP metabolic process | GO:0046034 | 32 | 7.5E-11 |
|  | ribonucleoside triphosphate metabolic process | GO:0009199 | 35 | 1.1E-10 |
|  | purine nucleoside triphosphate metabolic process | GO:0009144 | 34 | 1.5E-10 |
|  | nucleotide phosphorylation | GO:0046939 | 16 | 0.0056 |
|  | nucleoside triphosphate metabolic process | GO:0009141 | 37 | 0.0064 |
|  | purine nucleoside monophosphate metabolic process | GO:0009126 | 45 | 0.0149 |
|  | purine nucleotide metabolic process | GO:0006163 | 49 | 0.0244 |
|  | ribose phosphate metabolic process | GO:0019693 | 57 | 0.0261 |
|  | purine-containing compound metabolic process | GO:0072521 | 51 | 0.0269 |
|  | nucleoside monophosphate metabolic process | GO:0009123 | 48 | 0.0280 |
|  | ribonucleoside monophosphate metabolic process | GO:0009161 | 47 | 0.0310 |
|  | ribonucleotide metabolic process | GO:0009259 | 51 | 0.0348 |
|  | nucleoside diphosphate phosphorylation | GO:0006165 | 12 | 0.0354 |
| Amino acid metabolism | aromatic amino acid family metabolic process | GO:0009072 | 7 | 0.0627 |
|  | cellular amino acid metabolic process | GO:0006520 | 63 | 0.0512 |
| Lipid movement | lipid localization | GO:0010876 | 9 | 9.0E-11 |
|  | lipid transport | GO:0006869 | 9 | 2.2E-10 |
| Electron transport chain | electron transport chain | GO:0022900 | 12 | 0.0694 |
|  | ATP biosynthetic process | GO:0006754 | 10 | 0.0973 |
| Chemical homeostasis | chemical homeostasis | GO:0048878 | 14 | 0.0504 |
|  | cellular chemical homeostasis | GO:0055082 | 8 | 0.0720 |
| Cation transport | cation transmembrane transport | GO:0098655 | 35 | 0.0981 |
|  | cation transport | GO:0006812 | 42 | 0.0728 |
| Other | protein deubiquitination | GO:0016579 | 8 | 0.0999 |
|  | intra-Golgi vesicle-mediated transport | GO:0006891 | 5 | 0.0587 |

\* Benjamini Hochberg corrected enrichment p value

Many proteins related to cuticle formation did not map to GO terms, despite having significant BLAST hits to chitin-like proteins. To analyze how cuticle formation may be developmentally regulated, we manually retrieved all proteins with significant BLAST hits to chitinases, structural chitin and chitin binding proteins. Indeed, we observed stark differences in the types of chitinases and structural chitin that are utilized (**Figure 6B**). Young mites displayed a markedly different structural chitin profile than adult sons and daughters, which was different still compared to the armoured foundress. Relative expression levels for each protein can be found in **Supplementary Table 3**.

### Chromatin remodeling and positive regulation of transcription underlie sexual differentiation

*Varroa* follows the system of haplodiploid sex determination (i.e. females are diploid, males are haploid), but other than that very little is known about the mechanisms that contribute to sexual differentiation. To investigate this, we compared the proteins expressed in female (n = 9) and male (n = 6) mites and found 101 starkly differentially regulated proteins, providing a starting point on which to further investigate possible differentiation mechanisms (**Figure 7A**, **Table 1**). A disproportionately large fraction (over 80%) of the differentially regulated proteins were upregulated in the males. Investigating the 10 most significant proteins further, we found that only three had appreciable homology to sequences with known functions (**Figure 7B**) – uridine phosphorylase, histone lysine N-methyltransferase and heat-shock protein (HSP)83 – while the others either had no significant sequence similarities or the significant matches have not been functionally annotated. Despite this, functional enrichment analysis revealed that GO terms relating to chromatin remodeling, positive regulation of transcription as well as various metabolic processes were significantly enriched (**Table 3**, **Supplementary Table 2**). Intrigued by the prominent profile of HSP83, we further analyzed how the other HSPs are sexually regulated (**Figure 6C**). We found that there is a core group of three HSPs that are specific to the foundress, and another group of three HSPs are male-specific.

**Figure 6.**
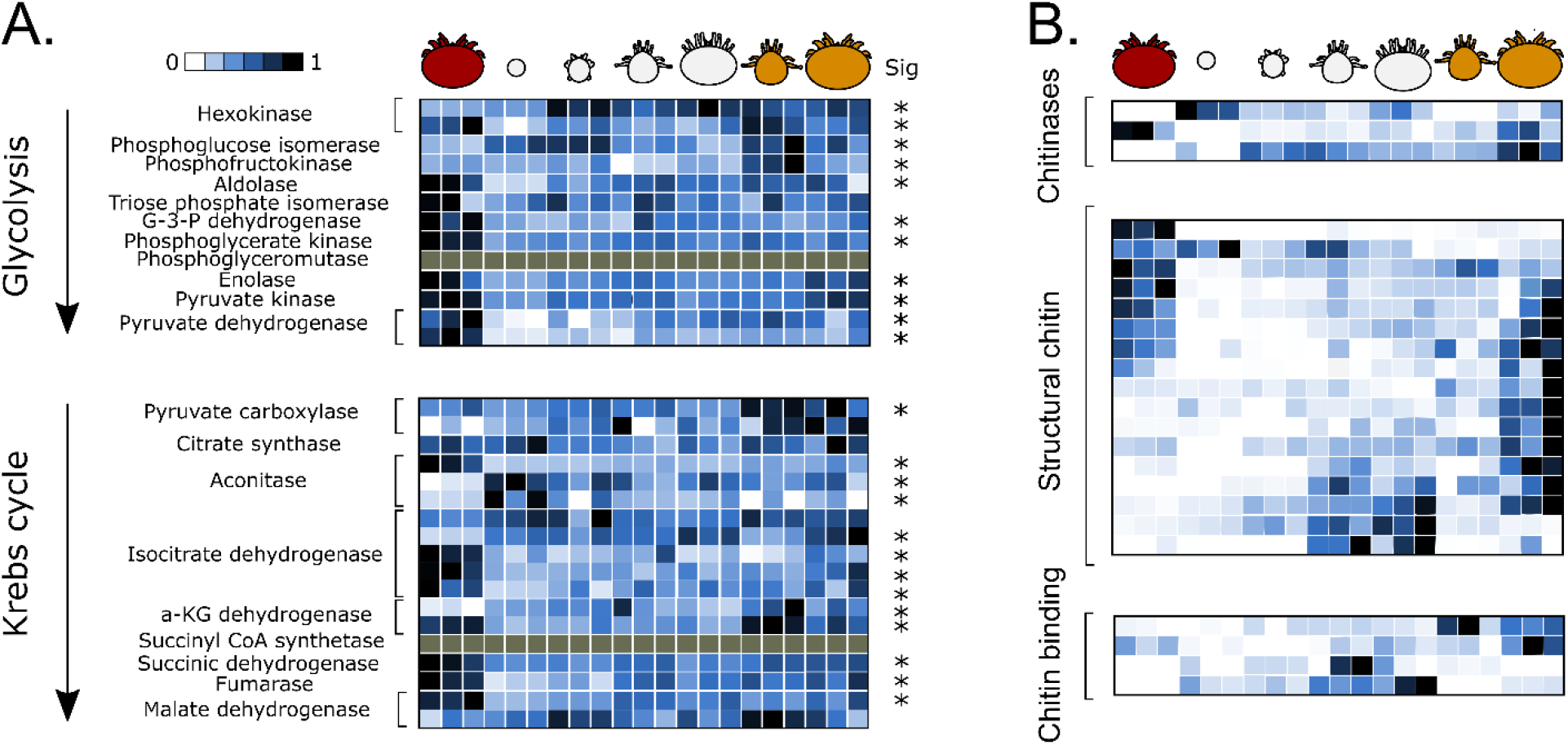
Analysis of carbohydrate metabolism enzymes and cuticle proteins. A. Relative expression of enzymes involved in carbohydrate metabolism. Bracketed rows indicate isoforms of enzymes catalyzing the same reaction (based on shared enzyme codes and having the enzyme in question as the best BLAST hit). Grey tiles indicate the protein was not observed. Rows indicated with an asterisk are significantly differentially expressed across developmental stages (from Figure 4A). G-3-P: glyceraldehyde 3-phosphate; a-KG: α-ketoglutarate. B. Relative expression of proteins related to chitin formation. Only the significantly differentially expressed proteins are shown.

**Table 3.** GO terms significantly enriched in sexually regulated proteins

| Description | Name | ID | # Genes | Corrected p value* |
| --- | --- | --- | --- | --- |
| Chromatin remodeling | DNA packaging | GO:0006323 | 7 | 0.0970 |
|  | DNA conformation change | GO:0071103 | 11 | 0.0698 |
|  | chromatin assembly or disassembly | GO:0006333 | 6 | 0.0833 |
| Transcription | positive regulation of gene expression | GO:0010628 | 6 | 0.0882 |
|  | positive regulation of transcription, DNA-templated | GO:0045893 | 5 | 0.0945 |
| Biosynthesis | positive regulation of biosynthetic process | GO:0009891 | 6 | 0.0950 |
|  | aromatic compound biosynthetic process | GO:0019438 | 70 | 0.0953 |
|  | organophosphate biosynthetic process | GO:0090407 | 43 | 0.0967 |
|  | nucleotide biosynthetic process | GO:0009165 | 37 | 0.0919 |
|  | glutamine family amino acid biosynthetic process | GO:0009084 | 6 | 0.0750 |
|  | organic acid biosynthetic process | GO:0016053 | 24 | 0.0798 |
| Metabolism/catabolism | cellular amino acid metabolic process | GO:0006520 | 64 | 0.0748 |
|  | alcohol metabolic process | GO:0006066 | 5 | 0.0838 |
|  | cellular nitrogen compound catabolic process | GO:0044270 | 13 | 0.0882 |
|  | organic cyclic compound catabolic process | GO:1901361 | 16 | 0.0882 |
|  | glycosyl compound metabolic process | GO:1901657 | 21 | 0.0882 |
|  | nucleoside metabolic process | GO:0009116 | 20 | 0.0907 |
|  | heterocycle catabolic process | GO:0046700 | 14 | 0.0937 |
|  | nucleobase-containing compound catabolic process | GO:0034655 | 9 | 0.0937 |
| Other | peptidyl-amino acid modification | GO:0018193 | 34 | 4.4E-10 |
|  | response to organic substance | GO:0010033 | 7 | 0.0682 |
|  | response to oxygen-containing compound | GO:1901700 | 5 | 0.0762 |
|  | protein folding | GO:0006457 | 48 | 0.0882 |

\* Benjamini Hochberg corrected enrichment p value

### Viral protein survey identifies DWV, BMLV and one ambiguous strain

*Varroa* is a vector for many damaging honey bee viruses and is a host to several itself (16, 17, 40-44), so we included all known *Varroa* and honey bee virus sequences in our protein search database. In total, we only identified three viruses – deformed wing virus (DWV), bee macula-like virus (BMLV) and one virus that is indistinguishable from *Varroa destructor* virus (VDV), DWV, or a hybrid of the two, although its polyprotein distinct from the aforementioned DWV strain. We will refer to the latter virus as VDV/DWV. Finding this number of different viruses is consistent with the findings of Erban *et al.* (2015), who identified three viruses within *Varroa* (DWV, *Varroa destructor* (Vd)MLV and acute bee paralysis virus). Interestingly, we found that DWV and VDV/DWV were significantly differentially regulated across developmental stages and showed distinctly different expression profiles (**Figure 8**). VDV/DWV displayed a marked decrease in expression in the egg and gradual increase through the remaining life stages, which is consistent with active replication in its host. DWV, however, exhibits a sharp increase in expression in adult daughters, but not in the other life stages. Two proteins corresponding to BMLV were detected: the coat protein (gi887503872) and P15 (gi887503873) – a protein translated from a short alternative reading frame at the 5’ end of the viral genome. P15 was detected in female and male deutonymphs and adult daughters, whereas the coat protein was detected only in female deutonymphs and daughters (**Table 4**).

**Figure 7.**
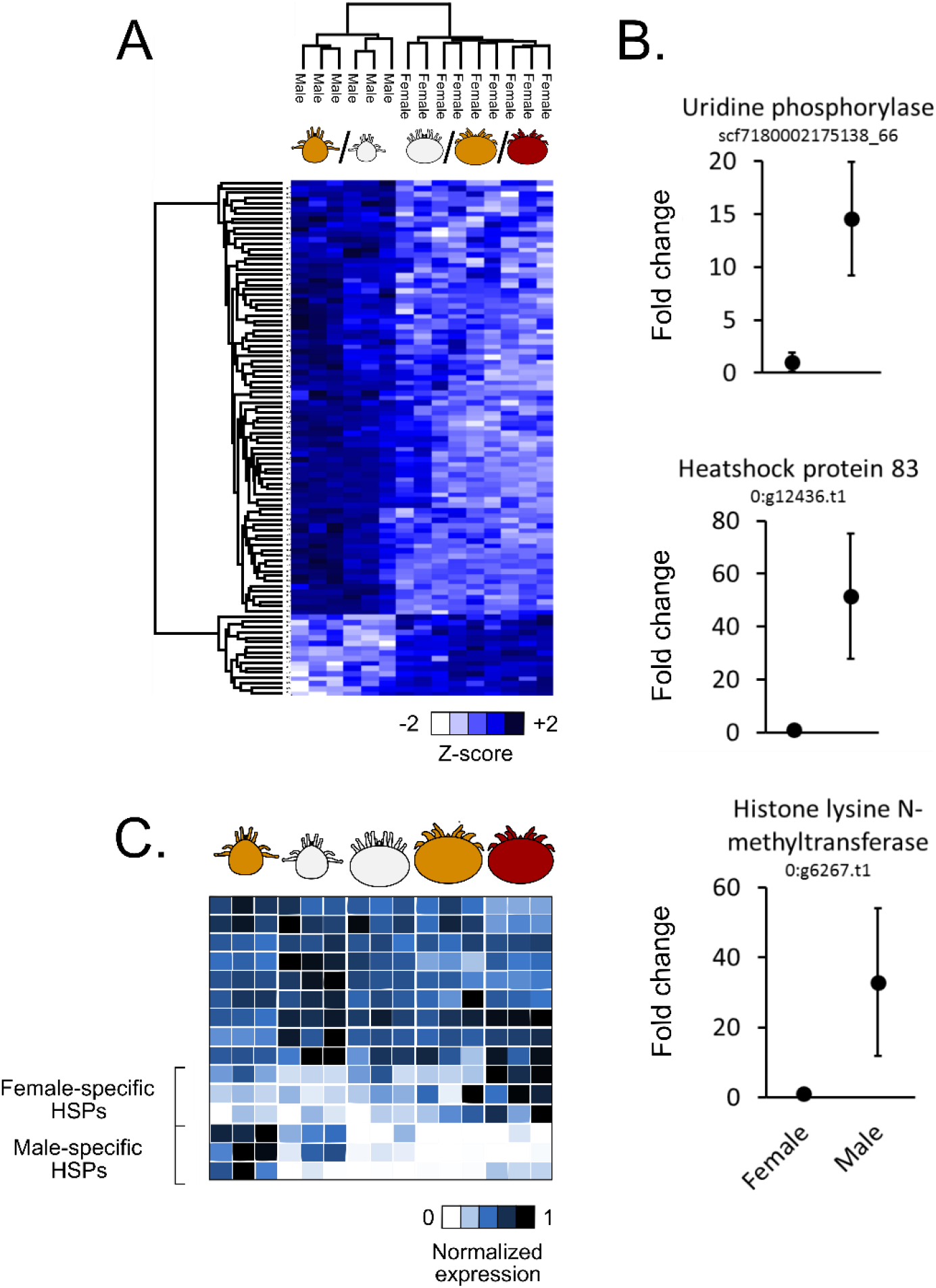
*Sexually regulated proteins in* Varroa. A) Heatmap showing differentially expressed proteins in male (n = 6) and female (n = 9) mites (Benjamini Hochberg-corrected FDR = 5%). Hierarchical clustering was performed using average Euclidian distance (300 clusters, maximum 10 clusters). B) The proteins with known functions among the top 10 differentially expressed. Fold change is normalized to the average expression in females. Error bars are standard deviation. C) Relative expression of heat-shock proteins (HSPs). Each row represents one HSP. Only significantly differentially expressed HSPs are shown.

**Table 4.** Abundances* of BMLV viral proteins

| Sample | Coat protein (BMLV) | P15 (BMLV) |
| --- | --- | --- |
| Daughter | 6.65 | 5.98 |
| Female deutonymph |  | 5.49 |
| Female deutonymph | 5.20 | 5.64 |
| Male deutonymph |  | 5.89 |
| Daughter | 6.55 | 6.13 |

\*log<sub>10</sub> transformed LFQ intensities

### Honey bee proteins are highly abundant in deutonymphs

*Varroa* mites feed on honey bee tissue, but how their diet might change during maturation is unknown. To examine how the honey bee proteins found within *Varroa* vary through development, we sorted out those proteins which were unequivocally honey bee-specific (i.e., the majority protein ID within the proteinGroups.txt output only contained honey bee accessions) and found that overall, the abundance of honey bee proteins increased drastically in the deutonymph stage (**Figure 9A**). We did not expect to observe any honey bee proteins in the egg samples, since the embryos are not consuming honey bee tissue; however, we still identified some, which is likely contamination from physical contact with the honey bee pupae. Interestingly, we observed a small group of honey bee proteins that were abundant mainly in the foundress (**Table 5**), and we hypothesized that this could arise if the foundress consumes honey bee tissues other than the hemolymph. To investigate this further, we determined the overlap between the observed honey bee proteins and a representative honey bee hemolymph proteome (provided by Hu *et al.* (45); PXD004467). We found that approximately 75% of honey bee proteins *Varroa* originated from the hemolymph regardless of life stage (**Figure 9B**). The origin of the remaining proteins is unknown. Normalized honey bee protein expression data for the deutonymphs and foundresses can be found in **Supplementary Table 4**.

**Figure 8.**
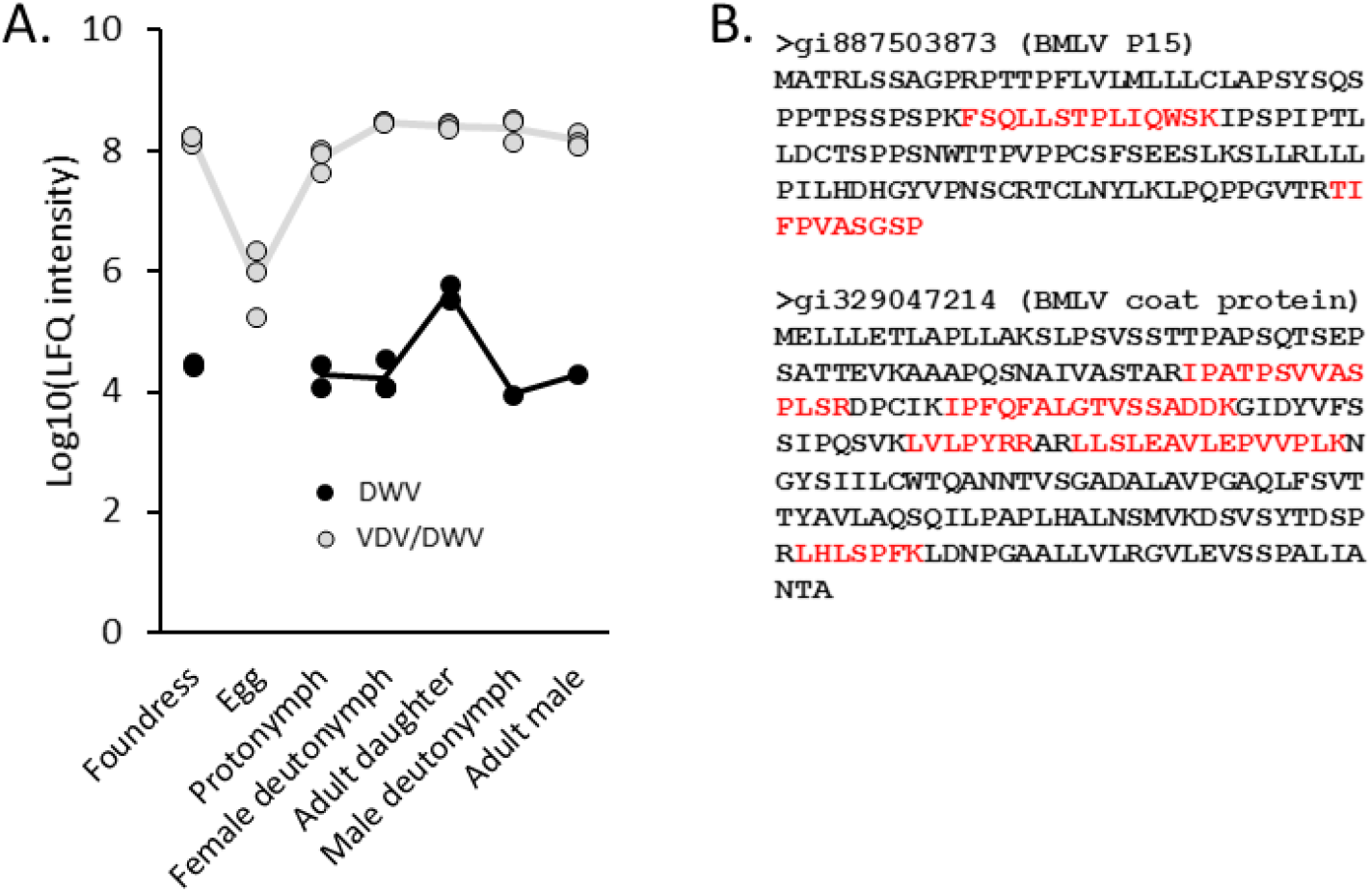
Survey of viral proteins. A) Deformed wing virus (DWV) and *Varroa destructor* virus (VDV)/DWV (a strain distinct from the aforementioned DWV) were significantly differentially expressed across developmental stages (ANOVA; Benjamini Hochberg-corrected 5% FDR). B) Bee macula-like virus (BMLV) proteins (1% protein and peptide FDR in MaxQuant). Red regions represent observed peptides.

**Figure 9.**
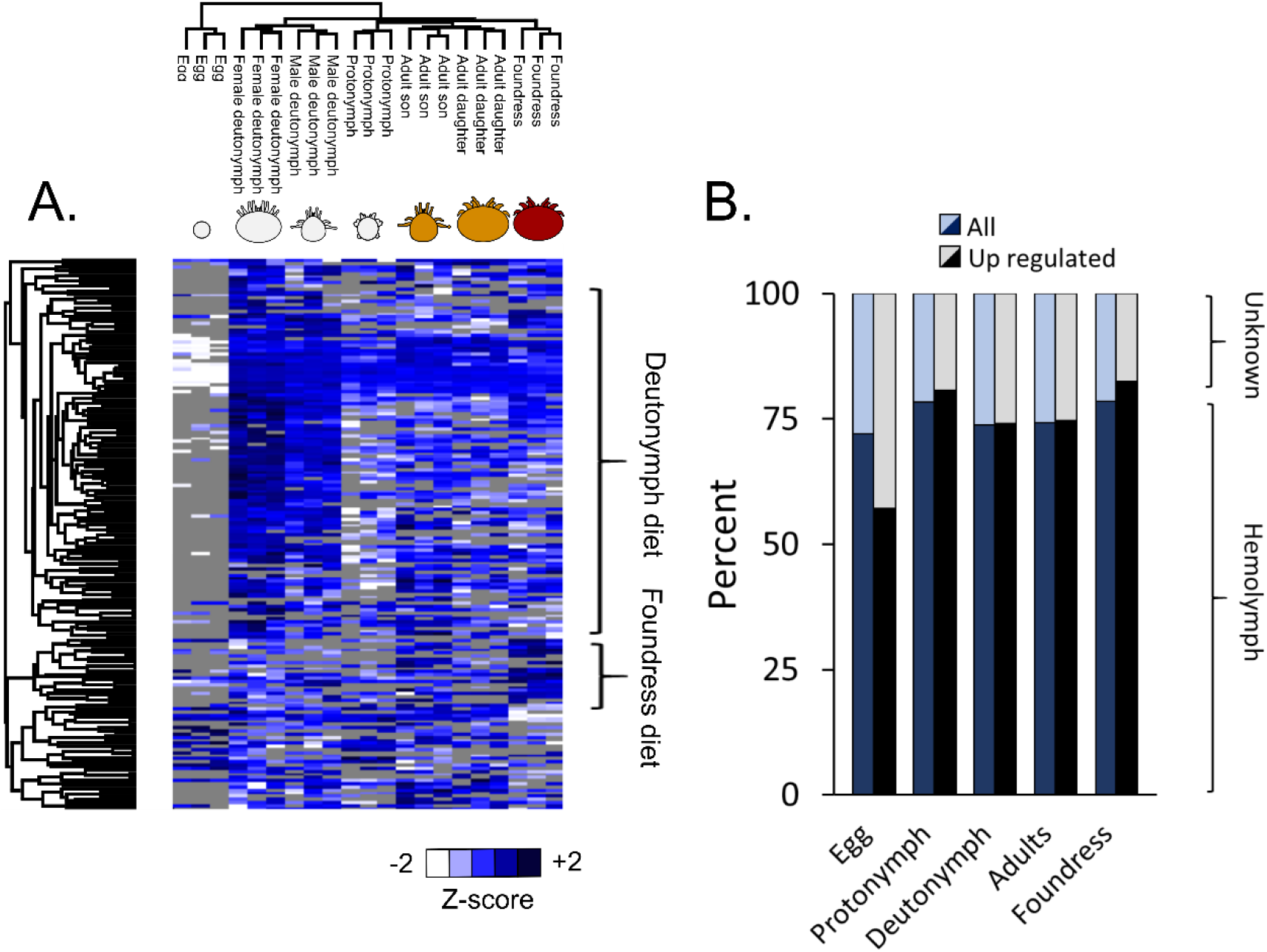
*Honey bee proteins which were differentially abundant throughout* Varroa *development*. A) Grey tiles are missing data. Differential abundance is based on Benjamini Hochberg-corrected 5% FDR. Hierarchical clustering was performed using average Euclidian distance (300 clusters, maximum 10 clusters). “Deutonymph diet” and “foundress diet” indicate distinguishable clusters of proteins which are highly abundant in those developmental stages. B) Fraction of bee proteins originating from the pupa hemolymph (dark bars) and unknown sources (light bars).

**Table 5.**
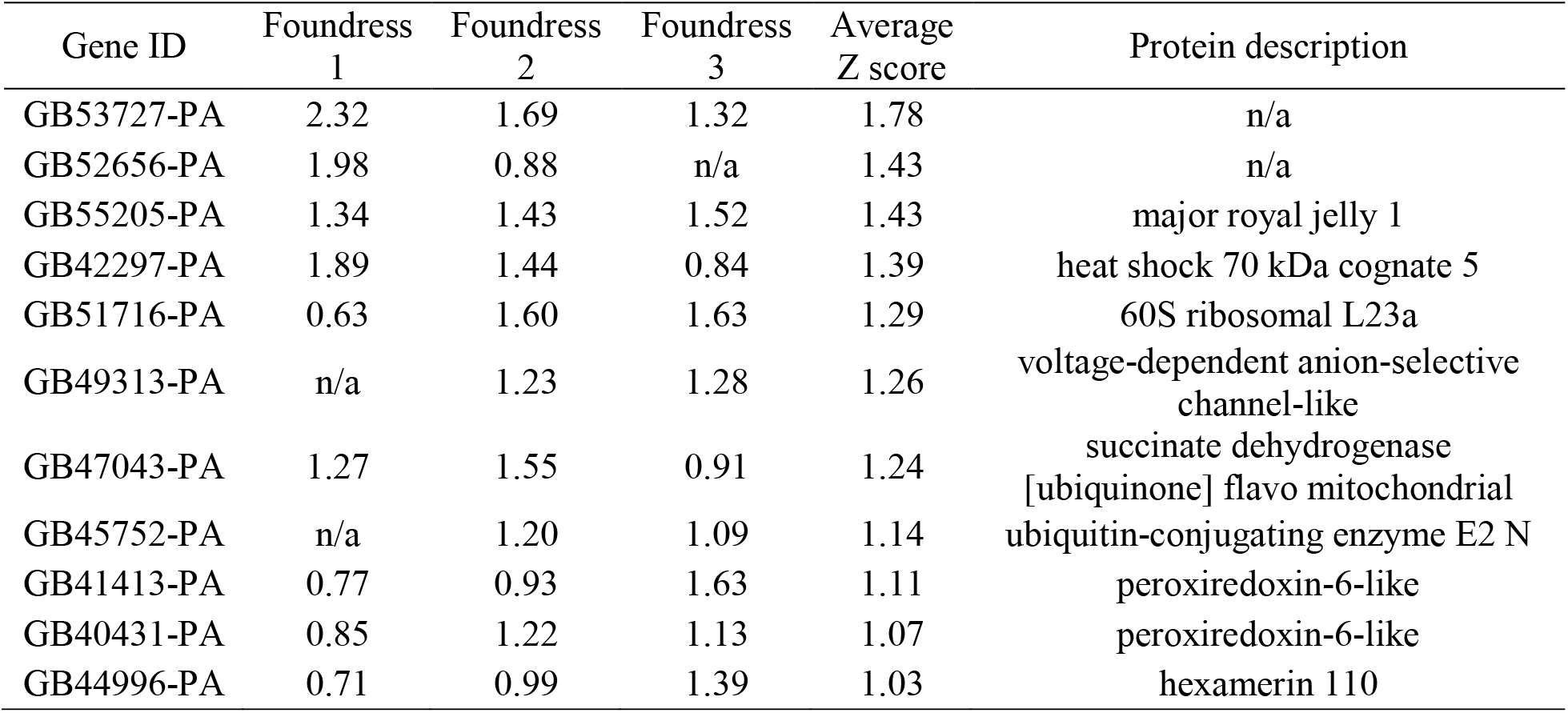
Honey bee proteins identified in the foundress with average Z score > 1

## Discussion

The work presented here provides a foundation to unravel the fundamentals of *Varroa* biology, including developmental transitions, sexual differentiation, diet, and host-virus interactions, as well as assisting with improving the genome annotation. Prior to this, there has been one published *Varroa* RNA-seq transcriptomics (24) and two proteomics studies, one of which – as far as we can tell – identified only virus and honey bee proteins and none from *Varroa* (15), and the other only analyzed foundresses (26). With almost 20,000 unique peptides identified, our study represents the deepest *Varroa* proteome to date. Overall, we identified 3,102 proteins, 2,626 of which were quantified by LFQ and incorporated into an interactive web-based *Varroa* proteome to serve as a community resource.

Genome sequencing is becoming relatively easy, but accurately annotating them is an arduous and imperfect process. The most common model organisms *(e.g. M. musculus, D. melanogaster, C. elegans, etc.)* have benefitted from decades of genetic research which has refined their genome annotations over time, resulting in highly reliable and accurate gene sets on which most global gene expression tools rely. Prior to the present body of work, expression studies in *Varroa* have been entirely limited to RNA analysis. This is the first time, to our knowledge, that protein expression of specific genes has been reported at all. Our data clearly show that the new *Varroa* gene annotation is far better than the provisional draft (**Figure 2**), but our proteogenomics initiative, which identified 1,464 unique unannotated peptides, suggests that there is still room for improvement. While some of these novel peptides simply harbour non-synonymous sequence polymorphisms, that itself is worth reporting and this information can be used to augment the protein databases used for mass spectrometry searches (46). Other peptides, however, clearly corresponded to exons of unannotated genes (**Figure 5C** and **5D**) which showed significant homology to vitellogenin. This observation, along with finding nothing unusual about the sequence properties of the newly identified coding regions (**Figure 4B – D**) led us to question why they were not already annotated.

The annotation process is not only influenced by the genome itself (chemical and physical properties, completeness, etc), but also by the quality of guiding transcript assemblies and a number of human-determined parameters (e.g. the annotation software employed, hard or soft repeat masking, splice site awareness, etc.), and availability of prior gene models (47, 48). Furthermore, some parameters may need to be altered on a species-by-species basis, but there is no inherent pathway for finding the optimal settings. Proteomics and RNA-seq data could serve as tools to not only confirm expression of predicted genes, but also to help define these parameters in the first place since the resulting protein and gene IDs are sensitive to database accuracy. The data we present here is all publicly available (PXD006072) and we urge future iterations of annotation refinement to take full advantage of this peptide evidence when developing new *Varroa* gene models.

In mass spectrometry-based proteomics, it is important that the protein database reflects the proteins that could be present in the sample. Since *Varroa* feeds on honey bee tissues and others have detected honey bee proteins in *Varroa* (15), we included honey bee proteins in the search database and found that 167 of them were significantly differentially abundant (**Figure 9A**). The eggs were largely lacking in honey bee proteins, which is in keeping with the developing embryos not yet being able to feed on wounded honey bee pupae. The presence of some honey bee proteins in the egg suggests these are contamination; however, the deutonymph stage of both sexes, which are actively feeding on hemolymph, showed a strong over-abundance of honey bee proteins. This suggests that the deutonymphs require large amounts of food, possibly to support energetically expensive developmental processes such as metamorphosis. As the mites reach adulthood, they appear to consume less honey bee material; however, the foundress displays an over-abundance of a small but unique group of honey bee proteins (**Table 5**). Regardless of developmental stage, approximately 75% of bee proteins overlap with a representative hemolymph proteome (**Figure 9B**), which does not support the notion that the *Varroa*’s food source switches from honey bee hemolymph to some other tissue.

Our analysis of developmentally-regulated proteins revealed some intriguing trends regarding the energetic demands throughout development (**Figure 6A**). The foundress had consistently high abundances of enzyme that participate in glycolysis and the citric acid cycle, which is possibly required to meet the energetic demands of producing and laying eggs. Some enzymes (e.g. hexokinase, α-ketoglutarate dehydrogenase, malate dehydrogenase and aconitase) have isoforms that are highly expressed in other developmental stages; however, the underlying purpose for expressing these particular isoforms is unclear. We speculate that many of the differences in metabolic processes are driven by the unique energetic requirements of metamorphosis, when energetically expensive morphological rearrangements must occur while the mite does not eat. During maturation, protonymph and deutonymph mites must transition from having a soft, translucent cuticle to acquiring a harder and more durable exoskeleton. The phoretic and foundress mites, in particular, have rigid armour to protect against injury by grooming honey bees and other environmental hazards. To investigate the possible mechanisms behind these transitions, we compared the expression profiles of significantly differentially expressed proteins that are related to cuticle development (chitin structural protein, chitinases and chitin binding proteins; **Figure 6B**). The egg contains large amounts of one chitinase and one chitin structural protein, which could be related to the breakdown of the egg case or the developing mite larva. The delicate protonymph, on the other hand, expresses only moderate amounts of a different chitinase and does not express appreciable amounts of any chitin structural proteins. Deutonymph stages, however, begin to display a specific profile of highly abundant structural proteins and chitin binding proteins, and from this point on there is a clear separation between male and female expression profiles. The male mite appears not to invest energy in forming a tough exoskeleton like the female does, which is consistent with the lack of environmental exposure during the male life cycle.

In our analysis of sexually regulated proteins, we found that chromatin remodelling and transcription activation were significantly enriched processes. Chromatin remodeling could be required to de-condense chromosomal regions which are highly expressed in males or females, and vice versa. Indeed, histone lysine N-methyltransferase was one of the most significant differentially expressed proteins, with approximately 30-fold higher levels in males compared to females (**Figure 7B**) and peptidyl-amino acid modification was the most significantly enriched biological process (**Table 3**). This kind of “on-off” regulation is consistent with our observation that numerous proteins were present in one sex and absent in the other (hence why it was advantageous to impute missing values for this analysis). Further exploring some of the most differentially expressed proteins, we found that HSP83 displayed the greatest fold change (~50-fold) out of those with known functions. This is consistent with previous observations in *Drosophila melanogaster,* which showed that this protein is critically important for spermatogenesis (49). Uridine phosphorylase – an important enzyme for uracil biosynthesis – was also one of the most significant proteins; however, it is unclear how it contributes to sexual differentiation. Since heat shock proteins are involved in transcription regulation, we further analyzed how their expression levels change through the life stages (**Figure 7C**). Indeed, there is a core group of HSPs that are specific to the foundress and another group that is specific to males, suggesting that these HSPs are involved in regulating the transcription of sex-specific genes.

Although we only observed proteins from three distinct viruses, this information provides some unexpected insights into viral replication strategies. Firstly, one viral polyprotein (which we call VDV/DWV) was indistinguishable between VDV, DWV and a recombinant VDV-1/DWV virus; however, it was distinct from another DWV strain we observed (**Figure 7A**) based on 12 unique peptides. The VDV/DWV polyprotein was above the 99^th^ percentile of most abundant proteins (out of all the 2,626 quantified). The extreme abundance of this pathogen-derived protein relative to *Varroa* proteins suggests that a significant portion of the mite’s energy resources are hijacked by the virus, possibly to reach excessively high titers. Previously, Zioni *et al.* (42) suggested that recombination between VDV-1 and DWV leads to a highly virulent strain that causes overt DWV-like symptoms. This hypothesis is consistent with our observation of a potential VDV/DWV recombinant polyprotein abundance that is approximately 10,000-fold higher than the DWV polyprotein. Of note, BMLV (Tymoviridae) is a recently discovered virus (50) and here we report the first expression confirmation, to our knowledge, of the protein encoded by the short 3’ overlapping reading frame (P15). For a non-structural viral protein, P15 displays remarkably high abundance (**Table 4**), even relative to the structural coat protein (which is normally by far the most abundant viral protein). This hints at it possibly also having a structural role in the BMLV architecture, but further studies will be required to elucidate it.

The work we present here represents a first-of-its-kind, high-resolution analysis of the *Varroa* proteome. With some 1,433 proteins that are differentially expressed, this data provides a first glimpse into the changes that take place during *Varroa* development. In addition, 101 strongly sexually regulated proteins provide clues for discovering the mechanisms behind sex determination and general dimorphism. We hope that the interactive web tool will maximize the utility of this information for the research community and will help generate further hypotheses for future experiments on this major honey bee pest.

## Footnotes

1 Jay Evans, personal communication (2017)

